# Human Skin Model from 15 GHz to 110 GHz

**DOI:** 10.1101/2025.05.26.652827

**Authors:** Andreas Christ, Adrian Aeschbacher, Bernadetta Tarigan, Ninad Chitnis, Arya Fallahi, Sven Kühn, Myles Capstick, Niels Kuster

## Abstract

In the recently revised guidelines for electromagnetic safety, basic restrictions expressed in terms of the absorbed power density (APD) at frequencies higher than 6 GHz were introduced. Testing for APD compliance of wireless devices requires experimental and numerical body models or phantoms that conservatively reproduce the absorption characteristics of human skin.

Previous studies of APD indicate that frequency-dependent impedance-matching effects are caused by the low-permittivity stratum corneum (SC) layer. The objective of this study is to complement previous work (Christ et al., 2020) and to develop dispersive dielectric models to represent reflection and absorption of electromagnetic fields at the surface of the skin across a frequency range up to 110 GHz.

The reflection coefficient of the skin of human volunteers was measured at frequencies of 15 to 43 GHz with open waveguide probes, complementing previous data from 45 to 110 GHz (Christ et al., 2020). The volunteers represented both sexes and different age groups and occupations; measurements were made at various regions of the body. The statistical analysis of the results show that the reflection coefficient follows a normal distribution in regions where the SC is relatively thin, which permits development of a conservative skin model that covers the 95^th^ percentile of the tested population. As expected, in regions where the SC is thicker, e.g., the palms, the reflection coefficient is not normally distributed, because the thickness of the SC depends on the mechanical stress and friction to which the hands are exposed during routine daily activities. There was no evidence of relevant differences due to sex, but there is evidence for a slight age-dependent difference.

The measured data – via fitting to the numerical model – allow the derivation of two-layer dielectric dispersive models that represent absorption and reflection at the surface of the skin with known uncertainty. The proposed models can be used to conservatively demonstrate compliance with the APD limits of wireless devices operating at frequencies of up to 110 GHz in any of the 5G and 6G bands defined.

**Highlights:**

- broadband evaluation of the skin reflection coefficient with open waveguide probes;
- corroboration of increased absorption of millimeter wave radiation in body regions with increased stratum corneum thickness;
- dispersive dielectric models representing reflection and absorption at the skin surface with known population coverage.

## 1 INTRODUCTION

In the most recent revisions of the exposure safety guidelines, the International Commission on Non-Ionizing Radiation Protection (ICNIRP) (ICNIRP, 2020) and the IEEE International Committee on Electromagnetic Safety (IEEE ICES) (IEEE C95.1, 2019) introduce new basic restrictions based on absorbed power density (APD) (ICNIRP, 2020) and epithelial power density (IEEE C95.1, 2019) averaged over an area of 4 cm^2^ for frequencies of 6-300 GHz, and 1 cm^2^ for frequencies above 30 GHz at the surface of the body. The objective of these restrictions is to limit the maximum temperature increase at the surface of the skin. The APD is a suitable quantity, because, at these frequencies, the penetration depth is small, and the incident electromagnetic (EM) power is predominantly absorbed in the surface layers.

Testing compliance of wireless devices in terms of the APD requires the use of phantoms that can adequately simulate reflection and penetration of EM fields by human skin and allow the APD to be determined with known uncertainty (IEC/IEEE DTR 63572, 2025; Chitnis et al., 2025). Similar requirements exist for phantoms used to test over-the-air (OTA) performance (*CTIA Test Plan for Mobile Station Over the Air Performance, Revision 3.9*, 2019) of wireless devices. The development of such phantoms is based on the frequency-dependent reflection/penetration parameters for any polarization of incident fields.

The most widely used dielectric parameters for skin tissues (Baumgartner et al., 2024) are derived from open coaxial probe (OCP) measurements of the human forearm skin (Gabriel et al., 1996b, 1996a; Baumgartner et al., 2024) at frequencies of up to 20 GHz. The OCP method provides accurate values for homogeneous materials but has serious limitations when used to measure layered structures like skin. A recent review of the dielectric properties of biological tissues (Sasaki et al., 2022) includes discussions of coaxial probes (Chahat et al., 2011; Zhekov et al., 2019), time-domain spectroscopy (Pickwell et al., 2004; Sasaki et al., 2017), and open-ended waveguide probes (Alekseev & Ziskin, 2007). Time-domain spectroscopy can be used only on excised tissue samples *in vitro*, whereas OCPs and waveguide probes can be applied for *in vivo* measurements. Open-ended waveguide probes measure the reflection coefficient of the fundamental transverse electric (TE_10_) mode and have already been applied to skin tissue (Alekseev & Ziskin, 2007; Alekseev et al., 2008; Christ et al., 2021).

The envelope of minimum and maximum absorption in the skin was evaluated by means of a greatly simplified stratified model of tissue layers of variable thicknesses (Christ et al., 2020), which demonstrated increased absorption of millimeter waves at frequencies greater than 20 GHz due to impedance matching effects in the stratum corneum (SC) layer. However, measurements of the skin of 37 volunteers made in different regions of the body confirm that the difference in the thickness of the SC of the palm compared to the other parts of the body results in considerable differences in reflection and transmission (Christ et al., 2021).

## 2 OBJECTIVES

The findings reported by (Christ et al., 2021) were limited to the frequency range of 40 to 110 GHz. The objective of that work was to develop phantoms for evaluations of the OTA performance of handheld and body-mounted devices operated at frequencies above 6 GHz. However, the findings were inadequate for the derivation of skin models to represent the user group with conservative coverage (typically > 90%) for demonstrating compliance with the APD exposure limits in the near-field of handheld and body-worn wireless devices (ICNIRP, 2020; IEEE C95.1, 2019).

The goals of this study are to extend the work of (Christ et al., 2021) to close the frequency gap between 15 GHz and 45 GHz and to derive dispersive dielectric models representing the reflection and absorption of human skin for different user-group coverage factors. In addition, the validity and accuracy of the applied measurement method were checked with the help of a homogeneous gel phantom with known dielectric parameters covered with low-permittivity layers of Teflon (polytetrafluoroethylene (PTFE)).

## 3 MATERIALS ANMETHODS

### 3.1 Instrumentation

WR42 and a WR28 open-ended rectangular waveguides with flanges from Eravant (Torrence CA, USA) were connected to an Anritsu (Atsugi, Kanagawa, Japan) model MS46131A vector network analyzer (VNA). The WR42 waveguide was used to analyze the 14–28 GHz frequency band, and the WR28 was used to analyze the 21–42 GHz band.

To ensure that the surface of the soft tissues, such as skin, is flat across the flange openings, pieces of rigid high-density low permittivity foam of 4.8 mm thickness were inserted into the flange openings. The dielectric parameters of the foam insert from Rohacell HF51 (Darmstadt, Germany) were measured over the of 5–67 GHz frequency range with the DAK-1.2E-TL2 dielectric probe kit for thin-layer materials from Schmid & Partner Engineering AG (Zurich, Switzerland). In addition, the measured data were compared to measurements made with the DAK-R resonator from Schmid & Partner Engineering AG at discrete frequencies of 10, 17, 26, 35, and 45 GHz determined as *ϵ*_r_ = 1.09 *±* 0.03 and tan *δ* = 2.68 *×* 10^−3^ *±* 3 *×* 10^−4^ (Schmid & Partner Engineering AG, 2025). To ensure mechanical stability during the measurements, the waveguides were fixed in a customized bracket. Figure 1 shows the measurement setup with the WR42 waveguide.

**FIGURE 1.**
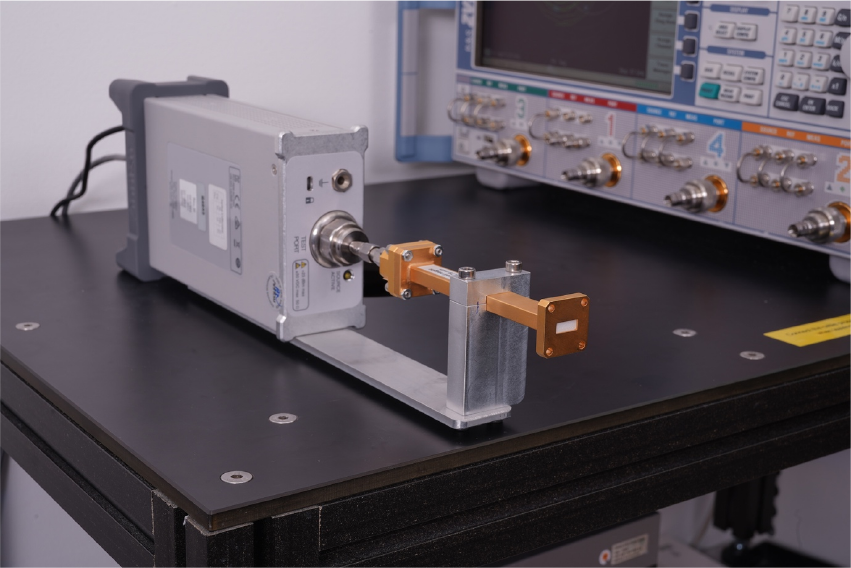
Measurement setup with the WR42 waveguide capped with rigid low permittivity foam and fixed with an aluminum bracket.

The WR42 and WR28 measurement setups were calibrated over their respective frequency ranges with a short, a *λ*/4 offset short, and with the STQ-TO-42-S1-CKIT1 and STQ-TO-28-S1-CKIT loads from Eravant. The low-loss foam and the adapter to connect the waveguide to the coaxial feed-line were included in the calibration. The reference plane for *S*_11_ is the interface between the waveguide flange and the measured sample

### 3.2 Volunteer Study

The measurements of the reflection coefficients *S*_11_ of the volunteers were carried out according to the protocol applied in the earlier study (Christ et al., 2021), summarized briefly here. The reflection coefficients of the skin of the 44 volunteers were measured at the locations shown in Figure 2. For each measurement, the waveguide was pressed against the skin of the volunteer. The distance between the waveguide flange and the measured surface, corresponding to the reference plane used for calibration, is considered to be zero.

**FIGURE 2.**
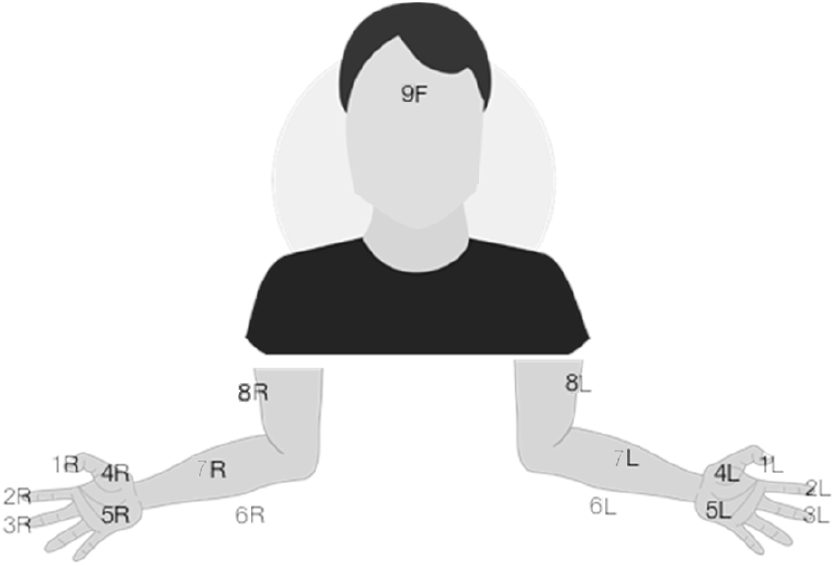
Locations on the hands, arms, and faces of the volunteers where *S*_11_ was measured, where F indicates forehead, L indicates left side, and R indicates right side.

Table 1 provides an overview of the age groups and sexes of the volunteers, and the *S*_11_ measurement sites on the hands, arms, and face (see (Figure 2) with thick vs. thin SC are listed in Table 2. Manual labor and hobbies that involve mechanical stress and friction on the hands are expected to be associated with an above-average increase of the thickness of the SC on the palms and fingers. The professions of 11 of the adult volunteers (over 20 years of age) in the study require manual labor; 25 are employed as office workers.

**TABLE 1.** Age and sex distribution of the volunteer participants. Data for WR15 and WR10 taken from (Christ et al., 2021).

| Age group | Age [years] | Sex | Number of subjects |  |
| --- | --- | --- | --- | --- |
|  |  |  | WR42, WR28 | WR15, WR10 |
| Child | 5–10 | female | - | 2 |
|  |  | male | - | 2 |
| Child | 10–15 | female | 3 | - |
|  |  | male | 5 | - |
| Adult | 20–40 | female | 6 | 5 |
|  |  | male | 6 | 8 |
| Adult | 41–60 | female | 7 | 7 |
|  |  | male | 8 | 8 |
| Elderly | 61–80 | female | 3 | 2 |
|  |  | male | 6 | 3 |
| Total |  |  | 44 | 37 |

**TABLE 2.** Measurement locations defined in Figure 2 with thin vs. thick SC layers. surface layer – referred to as Layer SC – and an inner layer – referred to as Layer D.

|  | Regions |
| --- | --- |
| Thick SC | 1L, 1R, 2L, 2R, 3L, 3R, 4L, 4R, 5L, 5R |
| Thin SC | 1F, 6R, 6L, 7L, 7R, 8L, 8R |

The *S*_11_ measurements were performed as follows:

- For each measurement site, 3 consecutive measurements of *S*_11_ were made and the mean was calculated.^*†*^
- Measured values of all regions with thick vs. thin SC (as listed in Table 2) were grouped and averaged.
- Mean and standard deviation values^*‡*^ for the two regions and the four waveguides (WR42, WR28, WR15, and WR10)^*§*^ were calculated, and the summary statistics for the different volunteer groups were evaluated.

### 3.3 Skin Model, Computational

#### Methods and Evaluation

As was reported in (Christ et al., 2021), absorption and reflection of the complex skin can be well approximated with a two-layer model stratified as a

The reflection coefficient of the skin tissue was evaluated computationally with the help of mode matching (MM) based on a code developed specifically for rectangular waveguides that terminate with a layered stratified dielectric. The method assumes that the waveguide flange is infinitely large and takes the fringing fields at the waveguide opening into consideration. As was observed in (Christ et al., 2021), the impact of the fringing fields on the reflection coefficient is relevant and must be taken into account, in particular for low permittivity dielectrics, such as the SC layer. A detailed description of the MM implementation for open waveguides can be found in (Chitnis et al., 2025). The implementation was validated with the help of measurements of a well-characterized coated dielectric as well as of simulations performed with the finite-difference time-domain (FDTD) method (Section 4).

The dielectric parameters of the material models of the skin (Section 5.3) and of the gel used for the validation measurements (Section 4) are represented by dispersive Cole-Cole and Debye models. The complex permittivity of the Cole-Cole model is given as

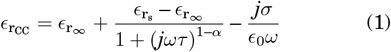

where 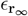 is the relative permittivity of the material when the frequency approaches infinity, 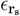 is static relative permittivity, *σ* is the static conductivity, *τ* is the relaxation time, and *α* is a parameter of value between 0 and 1. For *α* = 0, Eqn. (1) corresponds to the Debye dispersion model.

The parameters of Eqn. (1) were fitted to the measured reflection coefficients of the waveguides by means of sequential quadratic programming (SQP) in the implementation of GNU Octave (Eaton et al., 2022). The waveguide reflection coefficients for the optimization were calculated with MM (Section 3.3). The plane wave reflection coefficients were calculated analytically for the skin models on the basis of the fitted dispersion models. For the statistical analysis, R (version 4.0.3) and RStudio (version 1.3.1093) (R Core Team, 2017; RStudio Team, 2020) were applied.

## 4 VALIDATION OF THE APPLIED

### METHODS

The waveguide probe method (Section 3.1) and the calculation of the reflection coefficient by MM (Section 3.3) were validated with a homogenous lossy gel covered with PTFE having thicknesses of 50 *µ*m and 100 *µ*m. The dielectric properties are determined over a frequency range from 5 GHz to 67 GHz with the dielectric probe kit DAK 1.2E of Schmid & Partner Engineering AG. The uncertainty of the DAK 1.2E for this evaluation is < 0.2 dB.

The reflection coefficient of a tissue-simulating gel was measured with the waveguides WR42, WR28, WR15, and WR10. The numerical models of these waveguides, including their flanges and terminating with a layered dielectric were simulated by means of the FDTD method in Sim4Life Version 8.4 (Sim4Life.science, Zurich MedTech AG, Zurich, Switzerland). The implementation was verified according to (IEC/IEEE 62704-1, 2017).

The dielectric parameters of the gel can be represented by a Cole-Cole model as summarized in Table 3. The numerical results for the WR10 waveguide were obtained by extrapolation of the Cole-Cole model of the gel to frequencies up to 110 GHz. For the FDTD simulations, separate Debye models were fitted to this Cole-Cole model for the frequency ranges of each waveguide, and the waveguides were modeled with their flanges. Figure 3 shows the measured reflection coefficient compared to the results obtained with the MM and FDTD simulations. For all results, the maximum deviation is less than 0.5 dB.

**TABLE 3.** Dielectric models (Eqn. (1)) of the layered material used to validate the open waveguide probe setup.

| | d<br>mm | $\epsilon_{r\infty}$ | $\epsilon_{rs}$ | $\sigma$<br>S/m | $\tau$<br>ps | $\alpha$ |
| --- | --- | --- | --- | --- | --- | --- |
| Gel | >20 | 4.62 | 42.5 | 1.02 | 11.8 | 0.075 |
| PTFE | 0.05 and 0.1 | 2.1 | 2.1 | 0 | 0 | 0 |

**FIGURE 3.**
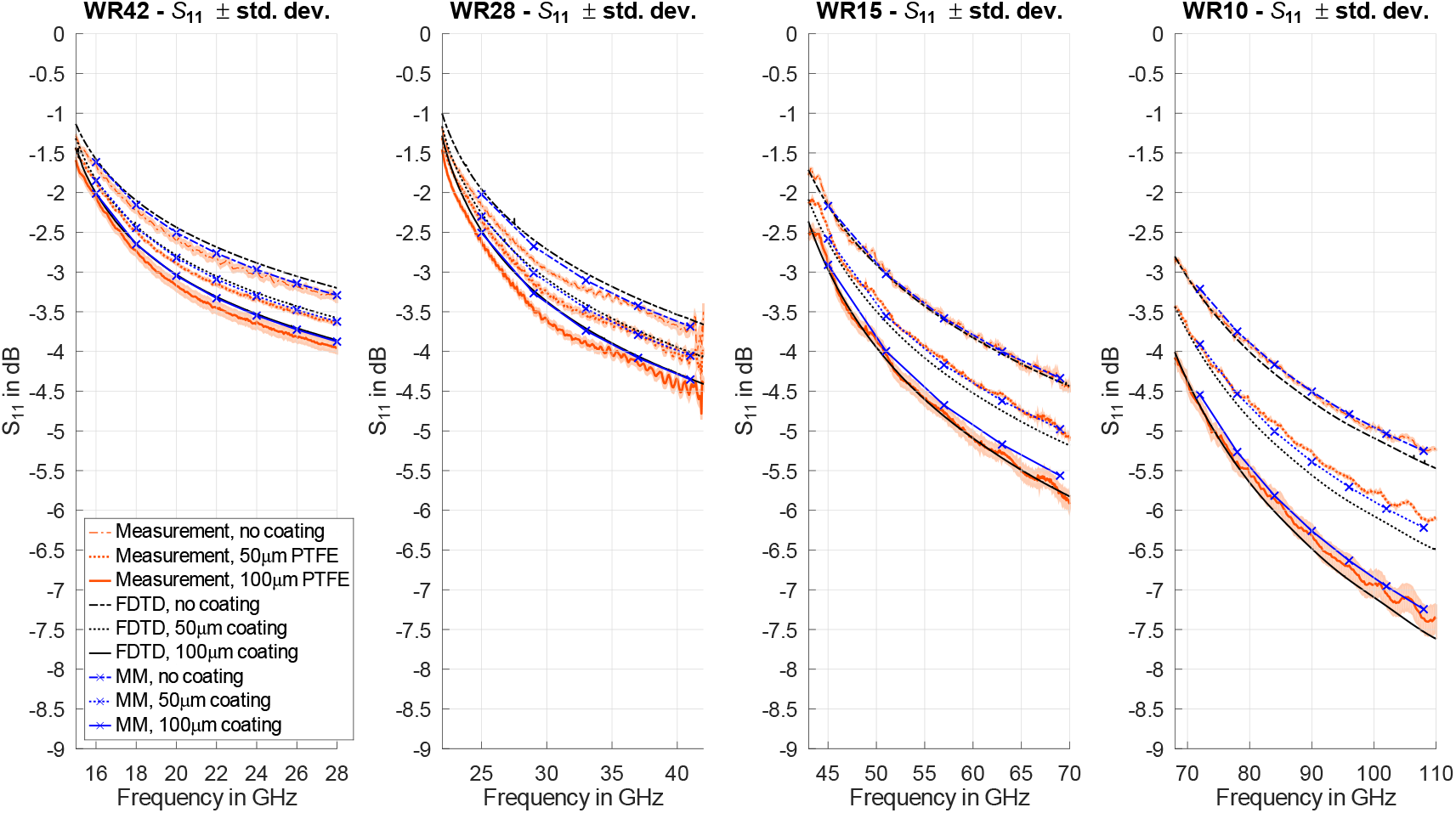
Measured reflection coefficients (mean of three measurements *±*1 standard deviation (shading)) of the TE_10_ mode of the tissue-simulating gel coated with two thicknesses of PTFE in front of the four waveguides used to cover the different frequency regions, compared to MM and FDTD simulations performed according to the dispersive layer model based on the parameters summarized in Table 3.

## 5 RESULTS

### 5.1 Statistical Evaluation of the SC Thickness

The distribution of *S*_11_ was tested for log-normal distribution at five distinct frequencies of the measurements with the WR42 and the WR28 waveguides. Table 4 summarizes the results of the Shapiro-Wilk test for regions with thick and thin SC (as indicated in Figure 2). A log-normal distribution could be demonstrated for both waveguides at all frequencies for thin SC only (*p* > 0.05). The measurement results for *S*_11_ in locations with thick SC show a 2-to 3-fold increase in variance, which was expected because SC thickness is a function of mechanical stress and friction of the palm and the fingers and is therefore not log-normally distributed.

**TABLE 4.** The Shapiro-Wilk tests on the difference of S_11_ (in dB) between thin and thick SC.

| Waveguide | Frequency | SC | $p$ |
| --- | --- | --- | --- |
| WR42 | 17.5 GHz | thin | 0.21 |
|  |  | thick | < 0.01 |
|  | 20 GHz | thin | 0.19 |
|  |  | thick | < 0.01 |
|  | 22.5 GHz | thin | 0.75 |
|  |  | thick | < 0.01 |
|  | 25 GHz | thin | 0.99 |
|  |  | thick | < 0.01 |
|  | 27.5 GHz | thin | 0.73 |
|  |  | thick | < 0.01 |
| WR28 | 24 GHz | thin | 0.61 |
|  |  | thick | < 0.01 |
|  | 28 GHz | thin | 0.33 |
|  |  | thick | < 0.01 |
|  | 32 GHz | thin | 0.16 |
|  |  | thick | < 0.01 |
|  | 36 GHz | thin | 0.40 |
|  |  | thick | < 0.01 |
|  | 40 GHz | thin | 0.45 |
|  |  | thick | < 0.01 |

**TABLE 5.** Results of the two-side paired *t*-tests on the difference in reflection coefficient between thin SC and thick SC Δ*S*_11_ and the lower and upper limits of the 95^th^ percentile confidence interval *l*_l_ and *l*_u_; *p* is always smaller than 2.2 *×* 10^−16^.

| Waveguide | Frequency | mean $\Delta S_{11}$ | $l_l$ | $l_u$ |
| --- | --- | --- | --- | --- |
| WR42 | 17.5 GHz | 0.47 | 0.43 | 0.52 |
|  | 20 GHz | 0.58 | 0.52 | 0.64 |
|  | 22.5 GHz | 0.69 | 0.62 | 0.76 |
|  | 25 GHz | 0.79 | 0.71 | 0.87 |
|  | 27.5 GHz | 0.91 | 0.82 | 1.00 |
| WR28 | 24 GHz | 0.49 | 0.44 | 0.54 |
|  | 28 GHz | 0.77 | 0.69 | 0.85 |
|  | 32 GHz | 1.05 | 0.95 | 1.16 |
|  | 36 GHz | 1.27 | 1.15 | 1.40 |
|  | 40 GHz | 1.53 | 1.40 | 1.67 |

This is illustrated in Figure 4, which shows several outliers that have *S*_11_ values of up to 0.5 dB lower than what would be expected for a log-normal distribution. This result differs from the findings reported by (Christ et al., 2021), where log-normal distribution could be shown for the difference between *S*_11_ at all frequencies of 60 GHz and above, with only one exception at 45 GHz. The different outcome can be explained by the more homogeneous group of volunteers in the earlier study.

**FIGURE 4.**
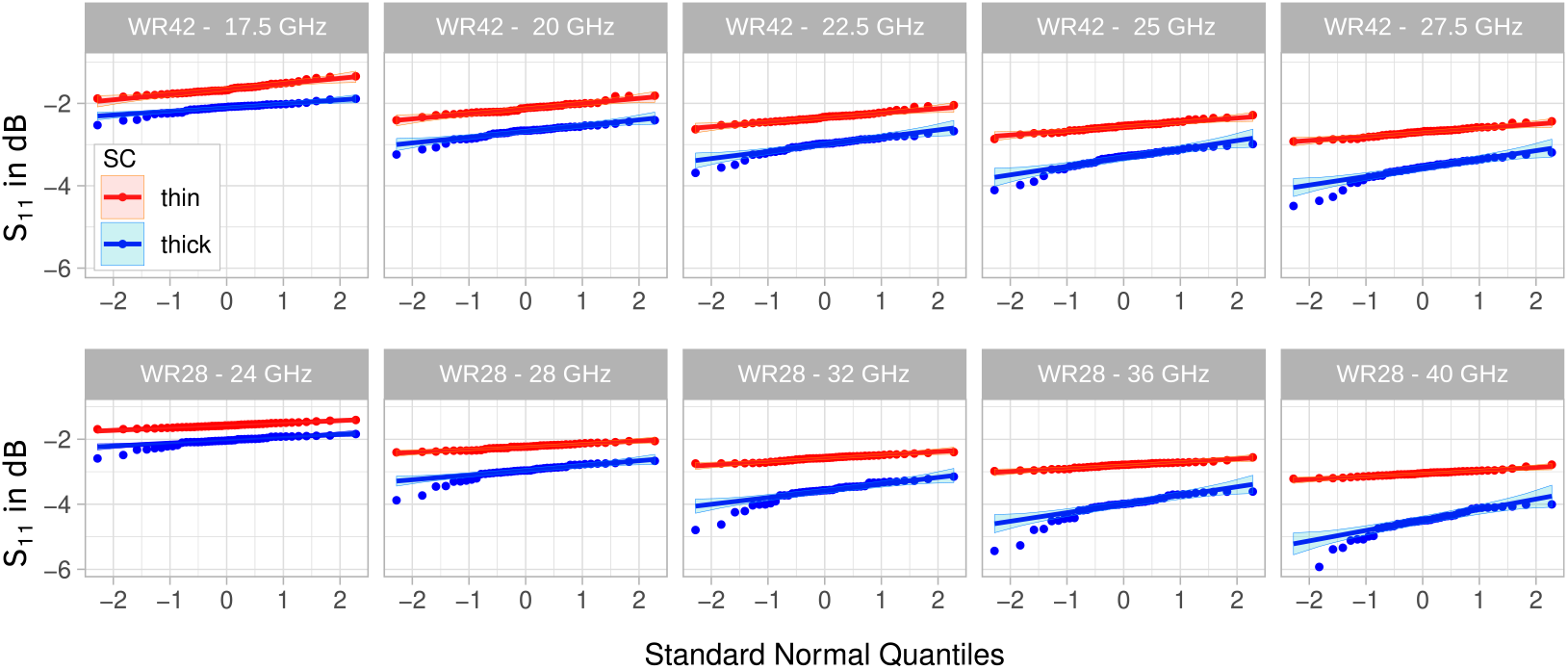
QQ plots of the *S*_11_ of all 44 volunteers measured with the WR42 (top row) and the WR28 (bottom row) waveguides at the body locations indicated in Figure 2 for thin (red) vs. thick SC (blue).

While normal distributions were assumed for body regions with thin SC only in the evaluations described in Sections 5.3 and 5.2 of this study, two-sided paired *t*-tests were carried out for the differences between S_11_ measurements of thick and thin SC, yielding a *p*-value of less than 2.2 *×* 10^−16^, which justifies the conclusion that the reflection coefficient of skin tissue can be classified as two distinct groups corresponding to thin and thick SC. Paired box plots and paired *t*-tests of the S_11_ measurements of thick and thin SC are shown in Figure 5.

**FIGURE 5.**
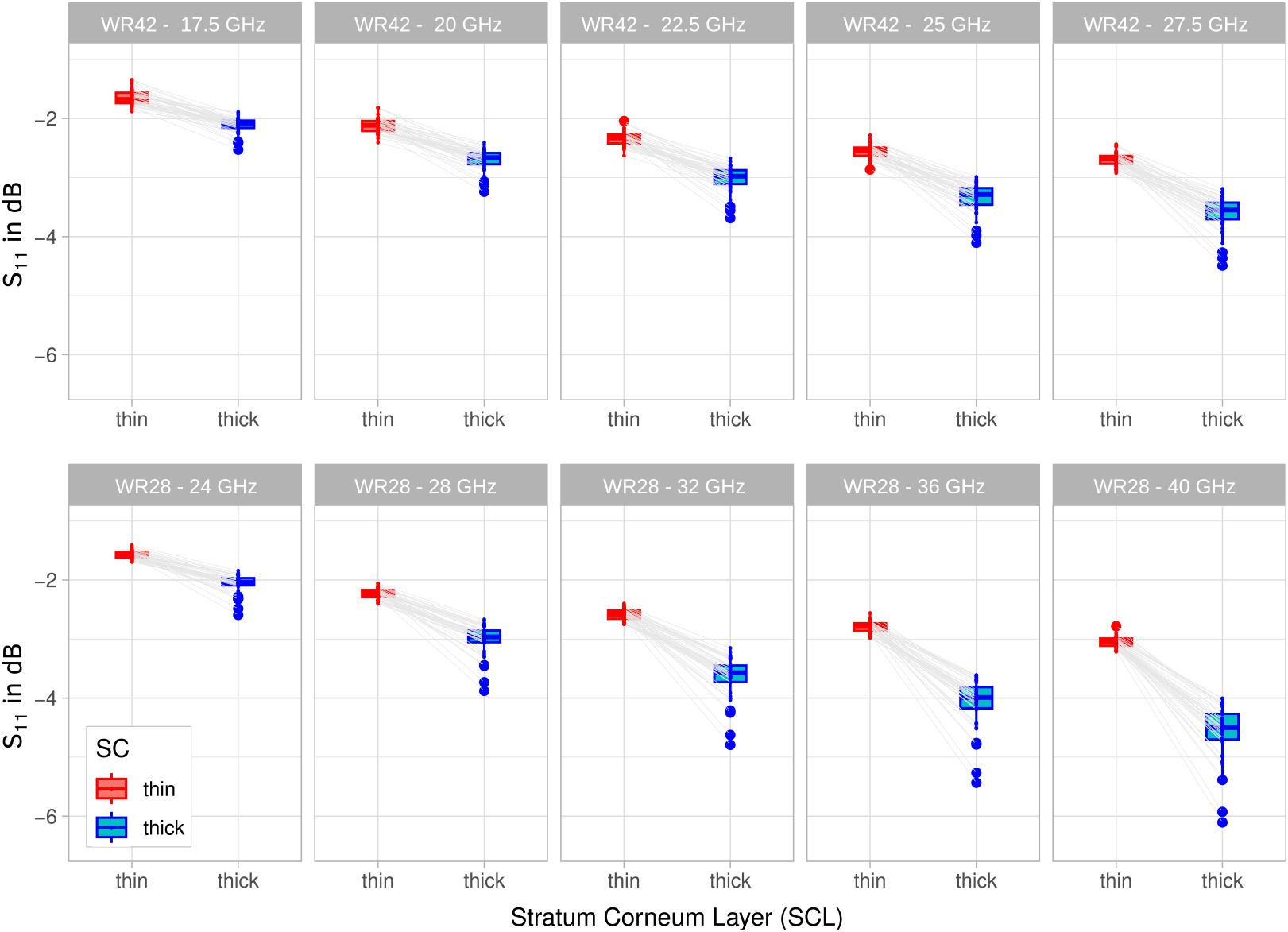
Paired box plots and paired *t*-tests (*p* < 2.2 *×* 10^−16^) for the *S*_11_ measurements performed at the body locations with thin (red) vs. thick SC (blue) layers of all 44 volunteers measured with the WR28 (top row) vs. the WR42 (bottom row) waveguide.

### 5.2 Statistics of the Volunteer Groups

In addition to the analysis of the SC thickness (Section 5.1), possible differences due to volunteer sex and age were evaluated. Weak evidence for sex-dependent differences had been observed previously in (Christ et al., 2021). The two-sample t-tests carried out on the measurement data in this study yielded *p*-values much larger than 0.05 (>0.5 in most cases), i.e., there was no confirmation of any evidence for sex-dependent differences. The Bartlett tests indicate equal variances (*p ≥* 0.30).

To test for age dependence, only the groups of young adults (20 – 40 years) and the elderly (60 – 80 years) were compared: the frequency dependence of the mean *S*_11_ was removed by normalizing the individual *S*_11_ to the respective mean value for each frequency. The two-sample *t*-tests indicate that the *p*-values are much smaller than 0.05, which indicates that age has an effect on the reflection coefficient of skin with thin SC. The observed decrease in the reflection coefficient for the elderly volunteers with respect to the young adult volunteers, however, is small (<0.2 dB) and need not be considered for the development of the dielectric skin models (Section 5.3). Again, the Bartlett tests indicate equal variances (*p ≥* 0.13).

### 5.3 Dielectric Skin Models for

#### Different Coverage Factors

The mean values and standard deviations of the reflection coefficients measured with the four waveguides are shown in Figure 6. The data for the WR15 and the WR10 waveguides were taken from (Christ et al., 2021). As already observed in Figure 5 and discussed in Section 5.1, the distributions of the reflection coefficients of thin and thick SC are clearly different.

**FIGURE 6.**
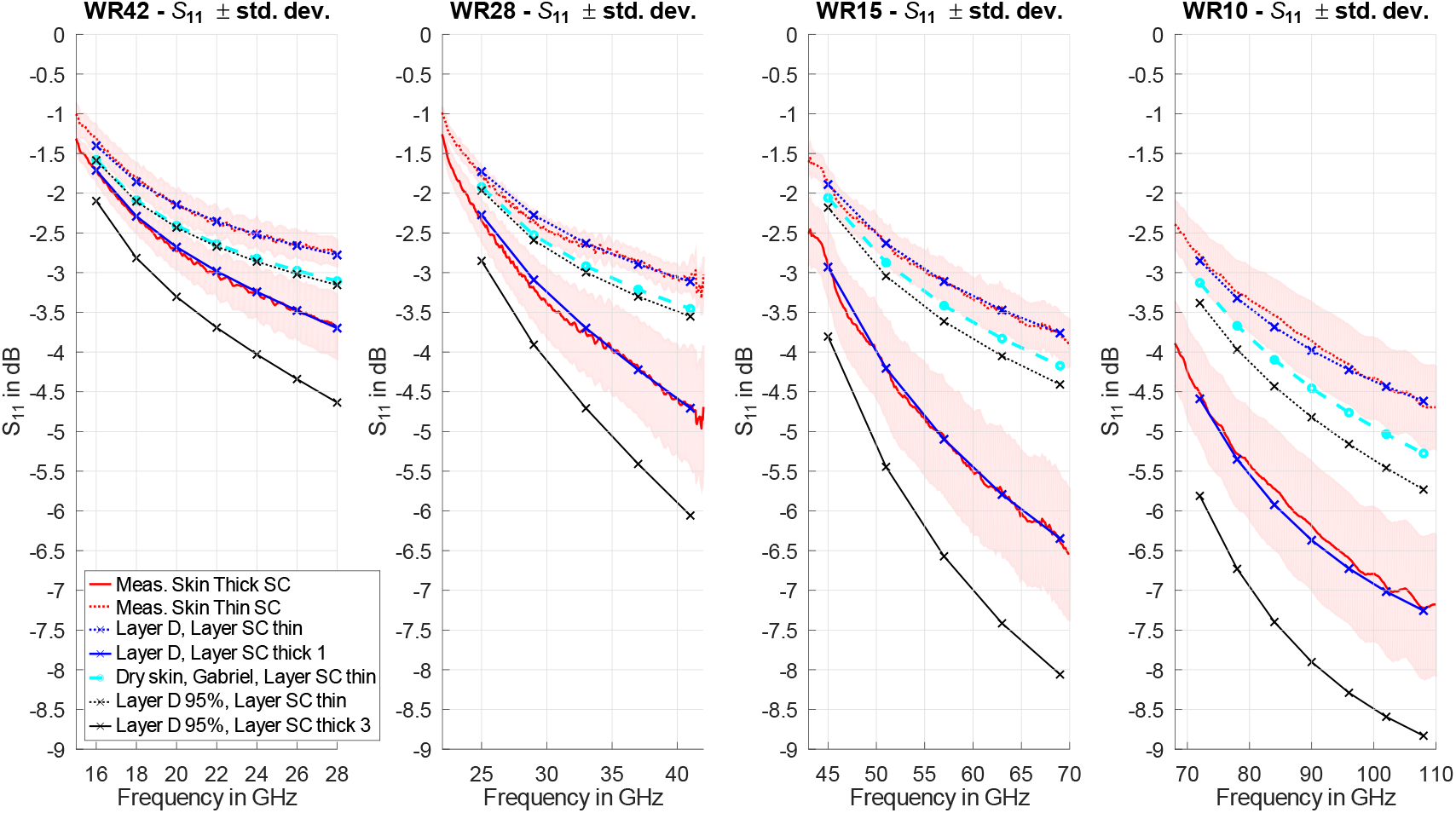
Measured reflection coefficients (mean of three measurements *±*1 standard deviation (shading)) of the TE_10_-mode measured with waveguides covering the different frequencies in front of locations with thick and thin SC and the optimized Debye models for Layer D and Layer SC. For comparison, the reflection coefficients of the dermis model of (Gabriel et al., 1996a) – evaluated with Layer SC – are included.

A Debye model for Layer D was fitted to the mean value of the reflection coefficient. For the fit, Layer SC with the dielectric parameters of (Ziskin et al., 2018) and a thickness of 20 *µ*m was used. This value is within in the range reported in (White et al., 1987;

Egawa et al., 2007) for body regions with thin SC. The results of the fit are shown in Figure 6, and the Debye parameters for Layer D are given in Table 6. Deviations of the fit from the measurement results are less than 0.1 dB. For a more conservative exposure estimate, additional models were fitted to the mean value of *S*_11_ minus one and two standard deviations. These correspond to the 68^th^ and 95^th^ percentiles of the coverage of the reflection coefficient and can therefore be used to assess the APD of locations with thin SC with known population coverage.

**TABLE 6.** Parameters of the Debye models (Eqn. (1) with *α* = 0) of the optimized dielectric models for layer thickness *d*. The parameters of Layer SC thin are taken from (Ziskin et al., 2018).

| | Comment | $d$<br>$\mu\text{m}$ | $\epsilon_{r\infty}$ | $\epsilon_{rs}$ | $\sigma$<br>S/m | $\tau$<br>ps |
| --- | --- | --- | --- | --- | --- | --- |
| Layer D | mean $S_{11}$ | ∞ | 7.88 | 47.0 | 5.19 | 8.35 |
| | 68 % coverage | ∞ | 5.06 | 37.0 | 7.62 | 8.19 |
| | 95 % coverage | ∞ | 2.98 | 32.8 | 6.94 | 8.76 |
| Layer SC | thin | 20 | 2.96 | 4.46 | 0 | 6.9 |
|  | thick 1 | 227 | 4.01 | 9.11 | 0 | 1.63 |
|  | thick 2 | 262 | 3.27 | 8.22 | 0 | 2.04 |
|  | thick 3 | 295 | 2.49 | 7.29 | 0 | 2.22 |

For the reflection coefficient of skin with thick SC, an alternative dispersive model for Layer SC, thick 1, was developed, for which a value of 227 *µ*m for the thickness was obtained from the fitting process. This value lies within the order of magnitude of the upper limit of the SC thickness reported, e.g., in (El Gammal et al., 1999; Egawa et al., 2007) for the palm and the fingers. Additional measurement results for Layer SC, thick 2 and thick 3, were fitted for the 68^th^ and the 95^th^ percentiles of the distribution of the measured reflection coefficient for thick SC.

The plane wave reflection coefficients of the skin models and for the dry skin model of (Gabriel et al., 1996a) with Layer SC of 20 *µ*m thickness were calculated analytically and are shown in Figure 7. For the 95^th^ percentile skin model with thin SC, the reflection coefficient ranges from –2.9 dB at 15 GHz to –6.0 dB at 110 GHz. The difference between the mean value and the more conservative 95^th^ percentile model ranges from 0.3 dB at 15 GHz to 1.0 dB at 110 GHz.

**FIGURE 7.**
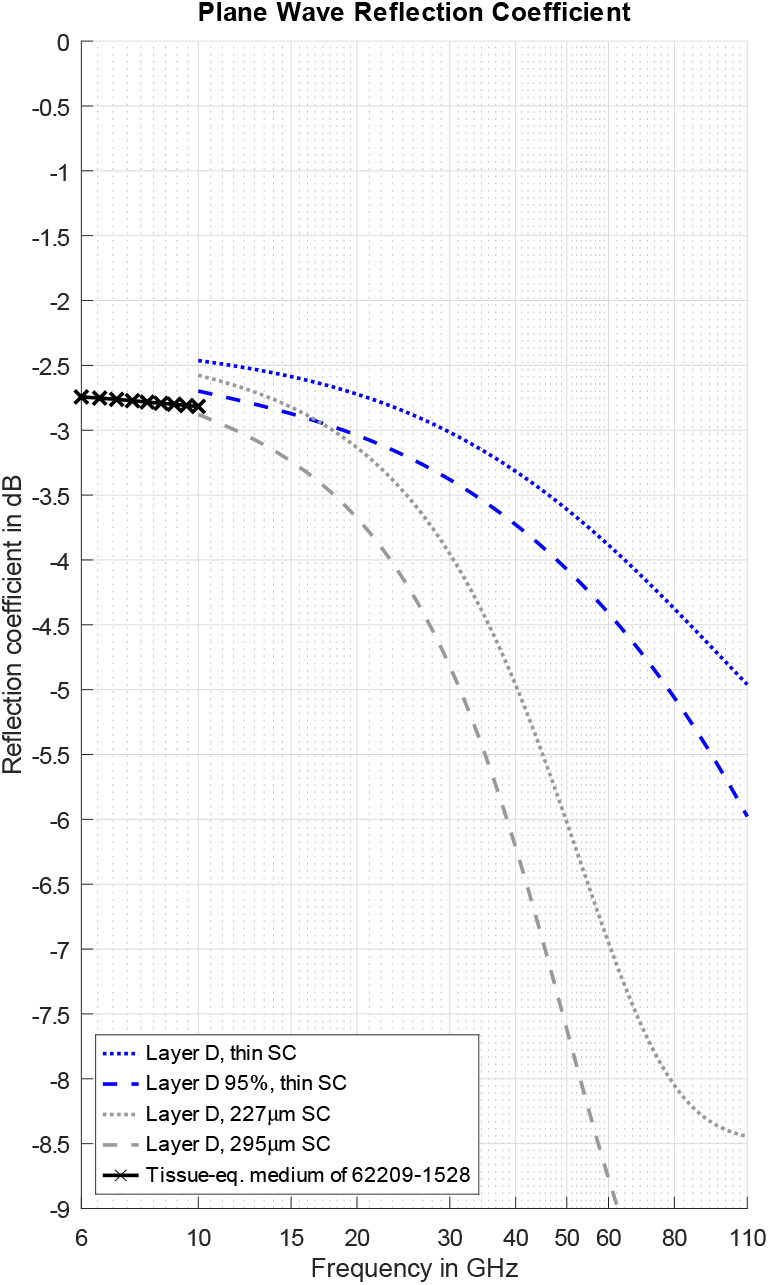
Plane wave reflection coefficients of the proposed conservative skin models for compliance testing of APD for a frequency range extending from 10 GHz to 110 GHz and compared to the tissue-equvalent medium (between 6 and 10 GHz) specified in (IEC/IEEE 62209-1528, 2020).

This model represents conservative skin absorption, whereas the model of (Christ et al., 2021), developed for OTA applications, reflects average values of the hands, the feet, and the body at frequencies >45 GHz derived from the same dataset, and offers comparable performance to existing phantoms below 6 GHz.

## 6 CONCLUSIONS

In this study, we investigated the reflection characteristics of human skin with the help of open waveguide measurements performed on 44 volunteers at locations with thick and thin SC, covering a frequency range of 15 – 43 GHz. The results confirm earlier observations – from a previous study for frequencies between 40 GHz and 110 GHz – ((Christ et al., 2021)) that absorption of EM energy is increased in regions with thick SC. The measurement results of both studies are sufficiently similar and could therefore be combined for evaluation. In detail, the findings can be summarized as follows:

- In the frequency range of 15 – 43 GHz, the difference in *S*_11_ could be shown to be statistically significant for the palms and the inner side of the fingers in comparison to the remaining body regions, which confirms the findings of previous studies.
- No evidence could be found for the dependence of the *S*_11_ for body regions with thin SC on the sex of the volunteers.
- When the youngest adult group (20 to 40 years) is compared to the elderly (60 to 80 years), there is evidence for a slight increase – less than 0.2 dB – in absorption of EM energy as a function of age.
- The combined results could be used to derive skin models to represent different coverage factors of the reflection and absorption characteristics for thin SC.
- Additional models for thick SC could also be derived as a function of the thickness of the SC layer.
- The plane wave reflection coefficient of skin with thin SC ranges from –2.9 dB at 15 GHz to –6.0 dB at 110 GHz for the 95^th^ percentile of the population.

Based on the extended frequency range covering all currently used 5G bands, a dielectric model for skin tissue is proposed that can be applied for the conservative assessment of the APD with a population coverage of *≈*95 %.

## Footnotes

† In total, 28 measurements that showed discontinuities in the slope of *S*_11_ were considered outliers and were not considered for further processing.

‡ See the discussion of the statistical distribution of the measurement results in Section 5.1.

§ The data for WR15 and WR10 were taken from (Christ et al., 2021).

## References

Alekseev, S., Radzievsky, A., Logani, M., & Ziskin, M. (2008). Millimeter wave dosimetry of human skin. Bioelectromagnetics: Journal of the Bioelectromagnetics Society, The Society for Physical Regulation in Biology and Medicine, The European Bioelectromagnetics Association, 29(1), 65–70.

Alekseev, S., & Ziskin, M. (2007). Human skin permittivity determined by millimeter wave reflection measurements. Bioelectromagnetics: Journal of the Bioelectromagnetics Society, The Society for Physical Regulation in Biology and Medicine, The European Bioelectromagnetics Association, 28(5), 331–339.

Baumgartner, C., Hasgall, P., Di Gennaro, F., Neufeld, E., Lloyd, B., Gosselin, M., … N, K. (2024). IT’IS database for thermal and electromagnetic parameters of biological tissues. http://www.itis.ethz.ch/database. Zeughausstr. 43, 8004 Zürich, Switzerland. Version 4.2, April 06, 2024.

Chahat, N., Zhadobov, M., Augustine, R., & Sauleau, R. (2011). Human skin permittivity models for millimetre-wave range. Electronics Letters, 47(7), 427–428.

Chitnis, N., Karimi, F., Kühn, S., Fallahi, A., Christ, A., & Kuster, N. (2025). Traceable assessment of the absorbed power density of body mounted devices at frequencies above 10 GHz. Bioelectromagnetics. submitted.

Christ, A., Aeschbacher, A., Rouholahnejad, F., Samaras, T., Tarigan, B., & Kuster, N. (2021). Reflection properties of the human skin from 40 to 110 GHz: a confirmation study. Bioelectromagnetics, 42(7), 562–574.

Christ, A., Samaras, T., Neufeld, E., & Kuster, N. (2020). RF-induced temperature increase in a stratified model of the skin for plane-wave exposure at 6–100 GHz. Radiation Protection Dosimetry, 188(3), 350–360.

CTIA Test plan for mobile station over the air performance, Revision 3.9. (2019). CTIA Wireless Association.

Eaton, J. W., Bateman, D., Hauberg, S., & Wehbring, R. (2022). GNU Octave version 7.3.0 manual: a high-level interactive language for numerical computations [Computer software manual]. Retrieved from https://www.gnu.org/software/octave/doc/v7.3.0/

Egawa, M., Hirao, T., & Takahashi, M. (2007). In vivo estimation of stratum corneum thickness from water concentration profiles obtained with Raman spectroscopy. Acta dermato-venereologica, 87(1), 4–8.

El Gammal, S., El Gammal, C., Kaspar, K., Pieck, C., Altmeyer, P., Vogt, M., & Ermert, H. (1999). Sonography of the skin at 100 MHz enables in vivo visualization of stratum corneum and viable epidermis in palmar skin and psoriatic plaquesy1. Journal of investigative dermatology, 113(5), 821–829.

Gabriel, S., Lau, R., & Gabriel, C. (1996a). The dielectric properties of biological tissues: III. Parametric models for the dielectric spectrum of tissues. Physics in Medicine & Biology, 41(11), 2271.

Gabriel, S., Lau, R., & Gabriel, C. (1996b). The dielectric properties of biological tissues: Ii. measurements in the frequency range 10 Hz to 20 GHz. Physics in Medicine & Biology, 41(11), 2251.

ICNIRP. (2020). International commission on non-ionizing radiation protection guidelines for limiting exposure to electromagnetic fields (100 kHz to 300 GHz). Health Physics, 118(5), 483–524.

IEC/IEEE 62209-1528. (2020). Measurement procedure for the assessment of specific absorption rate of human exposure to radio frequency fields from hand-held and body-worn wireless communication devices: Human models, instrumentation and procedures (Frequency range of 4 MHz to 10 GHz). Geneva, Switzerland: International Electrotechnical Commission (IEC), IEC Technical Committee 106.

IEC/IEEE 62704-1. (2017). Recommended practice for determining the spatial-peak specific absorption rate (SAR) in the human body due to wireless communications devices, 30 MHz-6 GHz — Part 1: General requirements for using the finite difference time domain (FDTD) method for SAR calculations. Geneva, Switzerland: International Electrotechnical Commission (IEC), IEC Technical Committee 106.

IEC/IEEE DTR 63572. (2025). Evaluation of absorbed power density related to human exposure to radio frequency fields from wireless communication devices operating between 6 GHz and 300 GHz. Geneva, Switzerland: International Electrotechnical Commission (IEC), IEC Technical Committee 106.

IEEE C95.1. (2019). Standard for safety levels with respect to human exposure to radio frequency electromagnetic fields, 0 Hz to 300 GHz. New York, USA: IEEE Standards Department.

Pickwell, E., Cole, B. E., Fitzgerald, A. J., Pepper, M., & Wallace, V. P. (2004). In vivo study of human skin using pulsed terahertz radiation. Physics in Medicine & Biology, 49(9), 1595.

R Core Team. (2017). R: A language and environment for statistical computing [Computer software manual]. R Foundation for Statistical Computing. Retrieved from https://www.R-project.org

RStudio Team. (2020). RStudio: Integrated development for R [Computer software manual]. R. RStudio, PBC. Retrieved from https://www.rstudio.com/

Sasaki, K., Mizuno, M., Wake, K., & Watanabe, S. (2017). Monte carlo simulations of skin exposure to electromagnetic field from 10 GHz to 1 THz. Physics in Medicine & Biology, 62(17), 6993.

Sasaki, K., Porter, E., Rashed, E. A., Farrugia, L., & Schmid, G. (2022). Measurement and image-based estimation of dielectric properties of biological tissues—past, present, and future—. Physics in Medicine & Biology, 67(14), 14TR01.

Schmid & Partner Engineering AG. (2025). Dielectric Database, Low-loss materials 2.0. https://speag.swiss/components/dielectric-database/low-loss-materials/., Zurich, Switzerland. Accessed: April 29th, 2025.

White, M., Jenkinson, D. M., & Lloyd, D. (1987). The effect of washing on the thickness of the stratum corneum in normal and atopic individuals. British Journal of Dermatology, 116(4), 525–530.

Zhekov, S. S., Franek, O., & Pedersen, G. F. (2019). Dielectric properties of human hand tissue for handheld devices testing. IEEE Access, 7, 61949–61959.

Ziskin, M. C., Alekseev, S. I., Foster, K. R., & Balzano, Q. (2018). Tissue models for rf exposure evaluation at frequencies above 6 ghz. Bioelectromagnetics, 39(3), 173–189.

